# Super-enhancer signature reveals key mechanisms associated with resistance to non-alcoholic steatohepatitis in humans with obesity

**DOI:** 10.1101/2021.08.20.457162

**Authors:** Yu-Han Hung, Ramja Sritharan, Marie-Claude Vohl, Olga Ilkayeva, Laurent Biertho, André Tchernof, Phillip J. White, Praveen Sethupathy

## Abstract

The molecular underpinnings of non-alcoholic steatohepatitis (NASH) development in patients are poorly understood. Active enhancer landscapes are known to determine cell states and behaviors. Super-enhancers, in particular, have helped reveal key disease drivers in several cancer types; however, they remain unexplored in human NASH. To define the enhancer signature of NASH-prone (NP) and NASH-resistant (NR) phenotypes in humans with obesity, we performed chromatin run-on sequencing (ChRO-seq) analysis on liver biopsies of individuals with obesity who were stratified into either NP or NR. We first demonstrated that NP and NR groups exhibit distinct active enhancer signatures. The subsequent identification of NP- and NR-specific super-enhancers revealed the specific genes that are likely the most critical for each of the phenotypes, including *HES1* for NP and *GATM* for NR. Integrative analysis with results from genome-wide association studies of NAFLD and related traits identified disease/trait-loci specific to NP or NR enhancers. Further analysis of the ChRO-seq data pointed to critical roles for serine/glycine metabolism in NASH resistance, which was corroborate by profiling of circulating amino acids in the same patients. Overall, the distinct enhancer signatures of human NP and NR phenotypes revealed key genes, pathways, and transcription factor networks that promote NASH development.

## Introduction

Non-alcoholic fatty liver disease (NAFLD) is the most common cause of chronic liver disease, affecting ∼25% of the population world-wide(1, 2). NAFLD is highly associated with many metabolic disorders including obesity, type 2 diabetes (T2D), and hyperlipidemia. Numerous risk factors have been identified, including body mass index (BMI), glycemic status, age, smoking, and specific genetic variants (such as rs738409 at the *PNPLA3* locus)(3–5). Clinically, NAFLD is characterized by abnormal accumulation of lipids in hepatocytes. In 10-20% of cases, NAFLD progresses to a more severe form known as non-alcoholic steatohepatitis (NASH)(2), which is marked by increased steatosis, hepatocyte ballooning, and lobular inflammation(6, 7), which sets the stage for the development of fibrosis, and in some cases eventually cirrhosis, hepatocellular carcinoma, or liver failure requiring transplantation(8). While NASH is the second leading cause, and is projected to soon be the primary cause, of liver transplantation in the United States(9, 10), there are currently no FDA-approved drugs for managing NASH(11). Many drugs that have shown promise for ameliorating NASH in animal models have failed in human clinical trials(12), which speaks to the urgent need for research with human tissue that aims to discover new potential therapeutic targets for NASH.

While gene expression profiles related to NAFLD progression have been reported previously in several studies(13–19), the underlying chromatin/transcriptional programming and the enhancer landscape that shapes gene expression in this disease remain unexplored in human patients. Regulatory elements (e.g., promoters and enhancers), which interact with transcription factors (TFs) to modulate activity of nearby genes, are critical for many fundamental biological processes including disease pathogenesis. The active enhancer landscape, in particular, very accurately distinguishes different cell states(20, 21). Furthermore, dense clusters of enhancers with remarkably high activity (known as super-enhancers) have been shown to mark genes that are critical for cell identity during development or cell behavior in disease pathogenesis(22, 23). While gene expression and/or regulatory landscapes in the context of liver fibrosis or NASH have been recently characterized in various hepatic cell populations sorted from mouse models(24–26), it remains completely unknown how the active enhancer/super-enhancer landscape and the associated TF networks are re-wired during NASH development in humans. Also, given that the current animal models of NAFLD and NASH may not fully recapitulate human NAFLD/NASH pathogenesis(24, 26, 27), we were motivated to leverage primary human liver biopsies to advance knowledge in the field.

The newest generation of nascent RNA sequencing technology developed to study chromatin regulatory dynamics(28), chromatin run-on sequencing (ChRO-seq), provides a powerful approach for characterizing the enhancer landscape and TF networks in human NASH. ChRO-seq enables sensitive measurements of transcription activity of gene bodies and transcriptional regulatory elements (TREs, such as promoters and enhancers) in one single assay, which greatly facilitates identification of enhancers and nearby associated transcriptionally-activated genes. Several key features of ChRO-seq distinguish it from other widely used approaches for studying gene regulation and TF networks. Notably, ChRO-seq detects TREs that are active, whereas the assay for transposase accessible chromatin sequencing (ATAC-seq) identifies all open chromatin loci, active or not. Furthermore, in contrast to ChRO-seq, open chromatin profiling methods do not provide quantitative information about gene transcription. Although transcriptional regulation and TF networks can be inferred from the promoter regions of genes that are differentially expressed in RNA-seq based studies, such a strategy harbors at least three major limitations. First, RNA-seq detects changes in genes at the steady state expression level, which is the result of both transcriptional and post-transcriptional layers of regulation. Second, promoters are less robust than enhancers for characterizing different cell states(20, 21). Third, RNA-seq does not provide any information at all about super-enhancers, which mark genes that are critical for the maintenance of cellular identity and behavior(22, 23). ChRO-seq ameliorates all of these limitations. Also, a major advantage of ChRO-seq relative to its previous iterations, known as GRO-seq(29) and PRO-seq(30), is that it can be applied to archived, frozen tissues, making it available for profiling human liver biopsy specimens. As an example of the utility of this method, recent studies have leveraged ChRO-seq to define transcriptional programs that underpin human cancer pathogenesis(28, 31) as well as gut development in a human organoid model(32).

In this study, we applied ChRO-seq on human liver biopsies from two well-matched groups of individuals with severe obesity that are discordant for the NASH phenotype to define the active enhancer signature, super-enhancer linked genes, and the candidate TF networks that are associated with NASH-prone (NP) and NASH-resistant (NR) livers.

## Results

### NASH resistant (NR) and NASH prone (NP) livers are associated with distinct profiles of gene transcription by Pol II

Many NAFLD studies in humans employ as the control group non-obese individuals who have lower BMI, more favorable metabolic profiles and/or younger age(18, 33–36). While such studies provide valuable information with translational implications, the presence of multiple confounders presents notable challenges for identifying factors specifically associated with NAFLD progression. Here we set out to study the enhancer landscape of NASH in humans using a comparison population that is matched for the major risk factors. Specifically, we first identified six individuals who have severe obesity (BMI > 45) without the presence of NASH. This cohort includes carriers of the *PNPLA3* risk allele as well as persons with impaired glucose tolerance (IGT) and type 2 diabetes (T2D) -- we term this group NASH-resistant (NR) (**Supplementary Table 1; Figure 1A-B**). We then identified eight individuals with NASH who were well-matched for BMI, sex, *PNPLA3* genotype, and glycemic status – we refer to this group as NASH-prone (NP) (**Supplementary Table 1; Figure 1A-B**).

**Figure 1.**
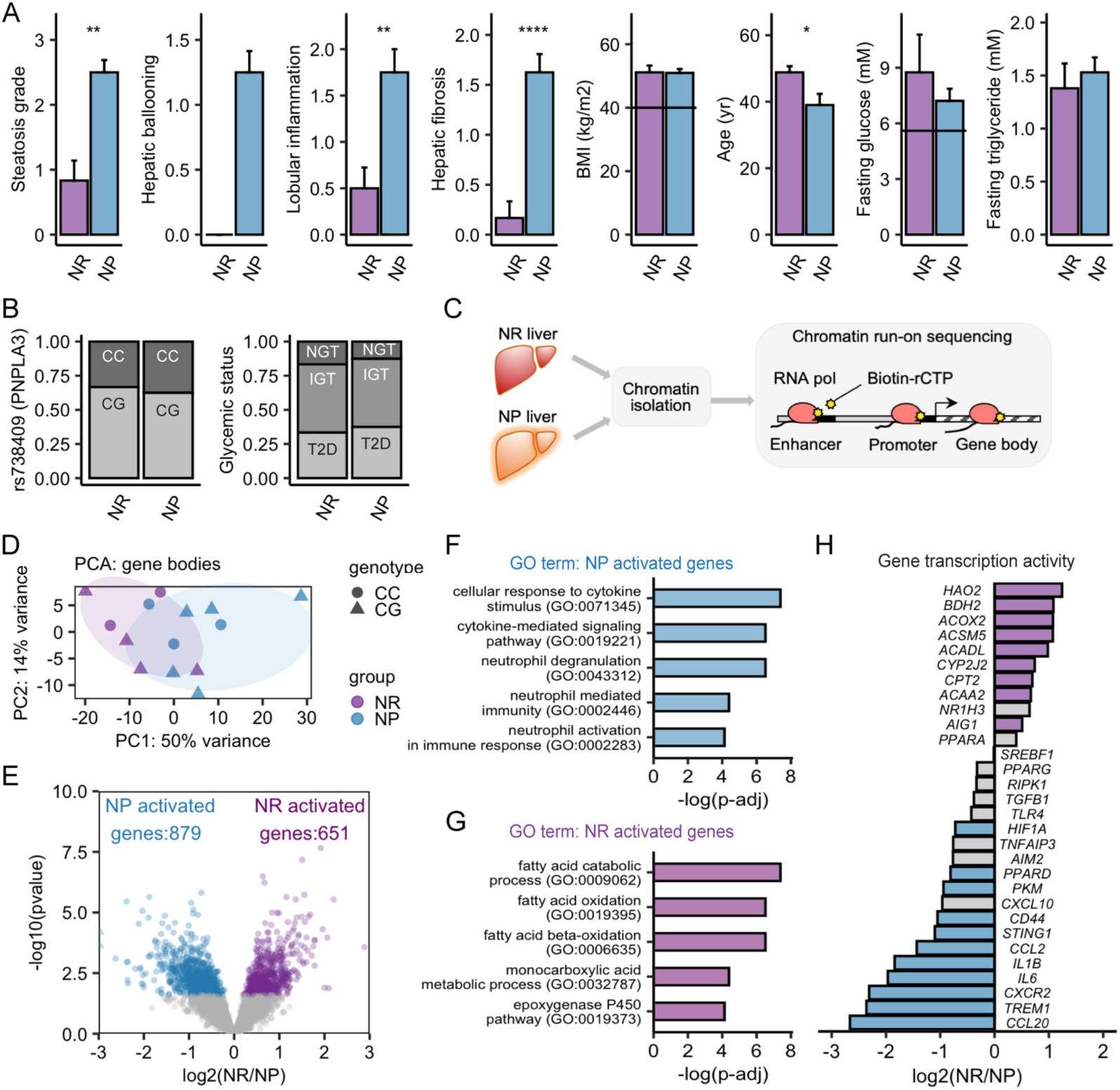
Distinct sets of genes are activated at the transcriptional level in human NASH-resistant (NR) and NASH-prone (NP) livers. (A) Clinical characterization of human subjects with NR and NP phenotype. In BMI panel, BMI above 40 (dashed line) is a general criterion for eligibility of bariatric surgery operation. In fasting glucose panel, values above 5.6 mM (dashed line) indicates hyperglycemia and risk for T2D. In HbA1c panel, values above 6.5 percent (dashed line) indicates poor glycemic control or T2D. * P < 0.05, ** P < 0.01, *** P < 0.001, **** P < 0.0001 by unpaired two sample T-test. See additional information in Supplementary Table 1. (B) Left: genotype of rs738409 at *PNPLA3* locus (CG, risk allele; CC, reference allele). Right: glycemic status of human participants with NR or NP phenotype. T2D, type 2 diabetes; IGT, impaired glucose tolerant; NGT, normal glucose tolerant. (C) Experimental scheme of chromatin run-on sequencing (ChRO-seq) profiling of primary liver biopsy samples from NR and NP subjects. (D) Principal component analysis (PCA) plot of ChRO-seq signal in gene bodies of NP and NR livers. Color denotes NP or NR phenotype. Shape denotes genotype of rs738409 at *PNPLA3* locus. (E) Volcano plot showing differentially transcribed genes between NP and NR livers. Genes with P < 0.05, adjusted P < 0.2, log2 foldchange > 0 (or < 0), average normalized counts > 100 were shown (Wald test; DESeq2). See complete gene list in Supplementary Table 3. (F-G) Pathway enrichment analysis of NP-activated genes (n = 879) and NR-activated genes (n = 651) based on Gene Ontology (GO) Biological Process 2018. (H) Histogram showing fold-changes in transcription activity for selected genes in NR compared to NP livers. Colored bars denote genes with P < 0.05, adjusted P < 0.2, log2 foldchange > 0 in NR compared to NP (Wald test; DESeq2). NASH-resistant (NR), n = 6; NASH-prone (NP), n = 8.

The livers from NP participants displayed the classic NASH phenotype, with higher steatosis levels and the presence of hepatocyte ballooning, lobular inflammation and tissue fibrosis (**Figure 1A**). In contrast, the livers from NR participants lacked hepatocyte ballooning and exhibited little-to-no lobular inflammation and fibrosis (**Figure 1A**). Individuals across both NP and NR groups exhibited very similar values of BMI, circulating lipids, glycemia and frequency of the risk allele of rs738409 at the *PNPLA3* locus (**Figure 1A-B**). Age is a known risk factor for NAFLD progression to NASH. In the present study, the confounding effects of age were minimized because the participants with NR phenotype were actually significantly older than those with NP phenotype (**Figure 1A**). We hypothesized that the molecular phenotype of NR livers holds important clues to limiting NASH progression. Numerous studies have demonstrated that disease-specific enhancers/super enhancers mark nearby genes that are critical for defining disease behavior(22, 23). However, the enhancer landscape of NASH progression in humans has not been characterized. To bridge this important knowledge gap, we sought to define the enhancer signatures of both the NR and NP phenotypes.

We first performed chromatin run-on sequencing (ChRO-seq) on the liver biopsies from individuals with severe obesity who were stratified into either the NP or NR group (**Figure 1C**), thereby generating information on the transcription level of genes, active promoters, and active enhancers in a single assay. Principal component analysis (PCA) of transcriptional signal at annotated gene bodies showed a modest separation between NR and NP livers (**Figure 1D**). Differential transcription analysis identified 879 and 651 genes that are more transcriptionally active in NP and NR, respectively (**Figure 1E**; p < 0.05, padj < 0.2, normalized counts > 100, log2FC < or > 0 by DESeq2). The genes that are more transcriptionally active in NP than NR (hereafter, referred to as NP-activated genes) are enriched in pathways involved in neutrophil activation, immunity, and cytokine responses (**Figure 1F**), such as *IL1B*(37), *IL6*(38), *CCL2* (encoding MCP-1)(39, 40), *CCL20*(*16*), *HIF1A*(41), *TREM1*(42), *PKM*(43), *CXCR2* (44), *CRP*(45, 46)*, IL32*(47), and *CD44*(48) (**Figure 1H**). The genes that are more transcriptionally active in NR than NP (hereafter, referred to as NR-activated genes) are enriched in pathways involved in fatty acid metabolism (*CPT2, ACADL, AIG1, ACAA2, ACSM5, ACOX2, BDH2, HAO2*) and xenobiotic processes (*CYP2J2*)(*49*)(**Figure 1G-H**). Overall, the transcriptionally activated genes and pathways in NP or NR are consistent with the literature(7), mostly based on studies of mouse models of NASH, describing candidate molecular players in NAFLD and/or NASH development.

### NR and NP livers have unique active enhancer signatures

ChRO-seq enables sensitive detection of active transcriptional regulatory elements (TREs), including promoters and enhancers. In this study, we defined a total of 136,863 active TREs across all samples, with an average size 397.1 bp (**Supplementary Figure 1A-B**). The identified active TREs were then classified as either promoter (n = 49,853; TREs overlapping −1000/+200 windows around transcriptional start sites) or enhancer (n = 87,010; TREs not overlapping −1000/+200 windows around transcriptional start sites). PCA of promoter and enhancer profiles showed that sample stratification was not driven by *PNPLA3* genotype (**Figure 2A**).

**Figure 2.**
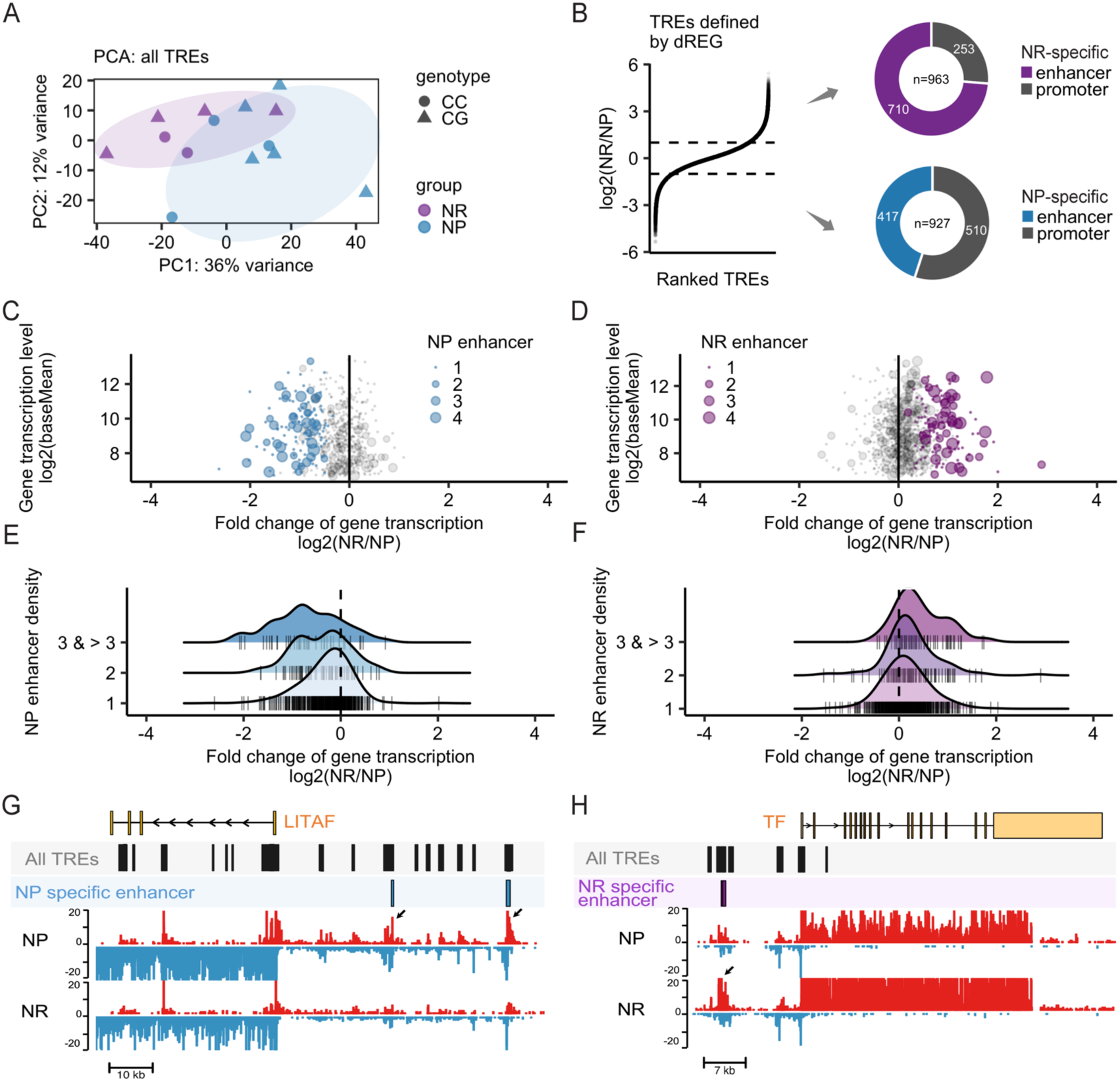
Active enhancer signatures of human NASH-resistant (NR) and NASH-prone (NP) livers. (A) Principal component analysis (PCA) of ChRO-seq defined transcriptional regulatory elements (TREs) in NP and NR livers. Color denotes NP or NR phenotype. Shape denotes genotype of rs738409 at *PNPLA3* locus. (B) Left: Ranked plot of all TREs, including both promoters and enhancers, based on foldchange of TRE activity in NR compared to NP livers. Dashed line denotes log2 foldchange at 1 and −1. Right: Donut plots showing numbers of differentially transcribed TREs between NP and NR livers. 927 TREs (417 enhancers and 510 promoters) are significantly active in NP livers. 963 TREs (710 enhancers and 253 promoters) are significantly activated in NR livers. P < 0.05, adjusted P < 0.2, log2 foldchange > 1 or < −1 in NR compared to NP by Wald test; DESeq2. See complete TRE list in Supplementary Table 3. (C-D) Ridgeline plot of genes (vertical tick marks) associated with one or more NP-specific (C) or NR-specific enhancers (D). Plots were generated for comparison across different enhancer-density groups. (E-F) Dot plot of genes associated with NP (E) or NR (F) specific enhancers. Size of the dot denotes enhancer density. Genes highlighted in colors are NP (blue) or NR (purple) activated genes (defined in Figure 1E); also see Supplementary Table 4. In (C-F), only genes with average normalized counts > 100 across all samples were shown. (H) Transcription activity (normalized ChRO-seq signal), all TREs, and NP-specific enhancers around locus of the *LITAF* gene are shown. (I) Transcription activity, all TREs, and NR-specific enhancers around locus of the *TF* gene are shown. NASH-resistant (NR), n = 6; NASH-prone (NP), n = 8.

Next, we performed DESeq2 analysis to define TREs that are strongly associated with NP or NR livers (**Figure 2C**; p < 0.05, padj < 0.2, log2FC < −1 or > 1 by DESeq2). We identified 927 and 963 TREs that are significantly activated in NP and NR, respectively (**Figure 2B**). Given that the samples exhibit more distinct enhancer profiles compared to promoter profiles (**Supplementary Figure 1C-F**), we focused the analyses on the subset of differentially active TREs that are enhancers, defined as NP-specific (n=417) and NR-specific enhancers (n=710) (**Figure 2B**). We next performed enhancer density analysis for each transcribed gene (average normalized counts > 100 across all the samples) by counting the number of NP- or NR-specific enhancers within a window of +/-100 kb around the annotated transcription start site (TSS). Several key observations were made in this analysis. First, there are only a small number of genes that are nearby to both NP- and NR-specific enhancers, indicating as expected that the sets of genes associated with NP- or NR-specific enhancers are largely mutually exclusive (**Supplementary Figure 1G**). Second, we found that genes that are associated with NP- or NR-specific enhancers are heavily skewed to be more transcriptionally active in NP or NR, respectively (**Figure 2E-F**). Third, genes with a greater density of NP- or NR-specific enhancers nearby exhibited greater transcriptional activation in NP and NR, respectively (**Figure 2C-D**). Together, these data indicate that the enhancer profiles are, indeed, shaping the transcriptional landscapes of each phenotype (NP/NR), which establishes the reliability of the enhancer signatures for further detailed analyses.

We identified 139 genes that are associated with 1 or more NP-specific enhancers and also exhibit significantly higher transcriptional activity in NP (**Figure 2E**), such as *APOBEC3A* and *LITAF*. The RNA editing enzyme APOBEC3A is known to promote pro-inflammatory macrophage activity(50). LITAF (**Figure 2G**) has been suggested to induce the pro-inflammatory phenotype associated with NASH in human livers(51). On the other hand, we identified 127 genes that are associated with 1 or more NR-specific enhancers and also exhibit significantly higher transcriptional activity in NR (**Figure 2F**), such as *HAO2* and *TF*. HAO2 has been reported to promote lipid catabolism in the context of renal carcinoma(52). Transferrin (*TF*) (**Figure 2H**) codes for an important liver-derived transport protein responsible for systemic iron delivery and metabolism, which has been linked to hepatic fibrosis in humans(53). By ChRO-seq analysis, we have defined NP and NR specific enhancers and their nearby genes.

### Enhancer clusters point to candidate driver genes of NR or NP phenotypes

Super enhancers, or enhancer hotspots, are hyper-active dense clusters of individual enhancers. Accumulating evidence shows that enhancer hotspots serve to control the transcription of genes especially critical for cell fate identity during development and cell behavior in disease progression(22). To identify enhancer regions, stitched enhancer regions, and enhancer hotspots in NP, we used an algorithm that we have recently developed(31, 32) (Methods; **Figure 3A**), which itself is adapted from a previous pipeline for super enhancer detection(23).

**Figure 3.**
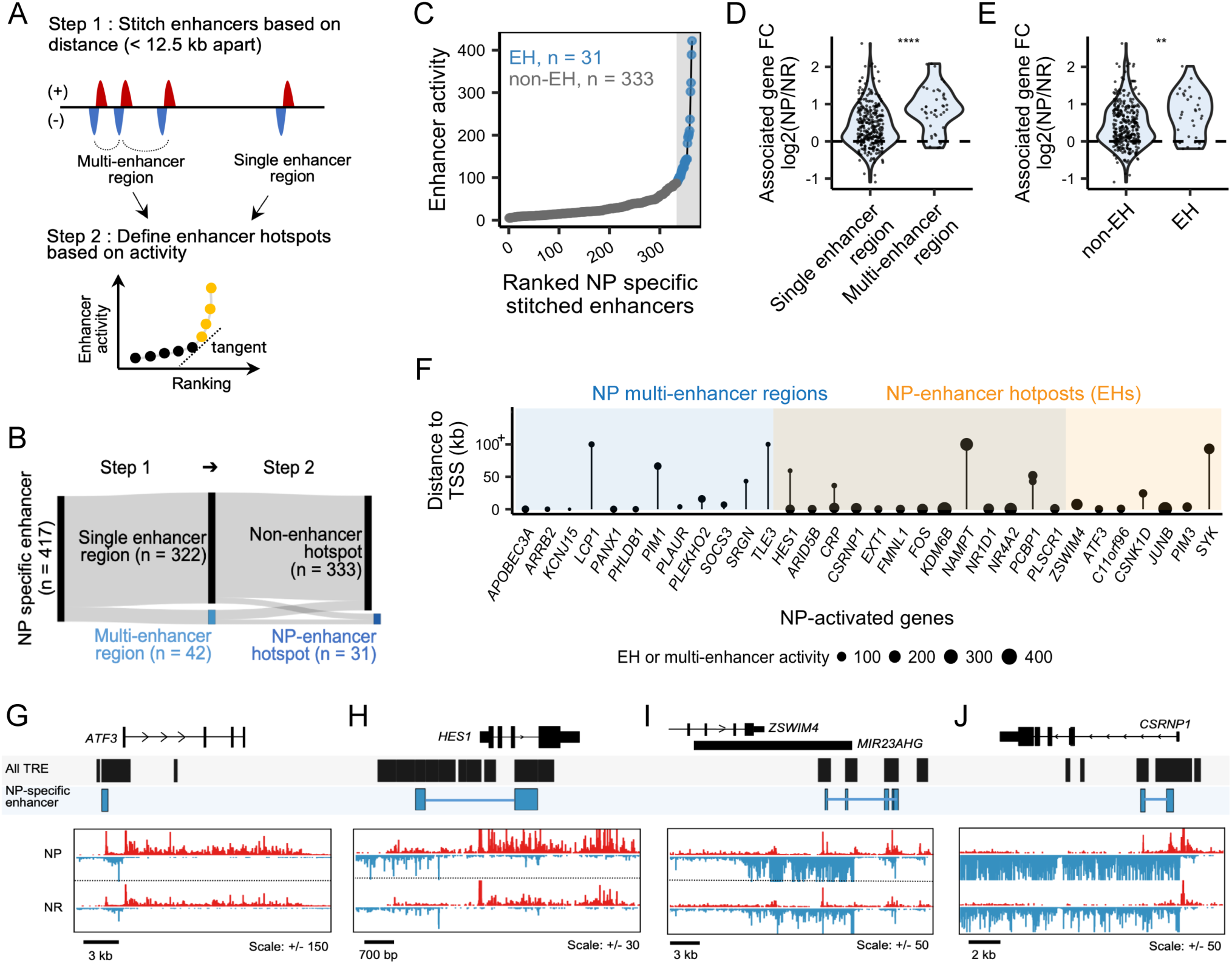
Identification of multi-enhancer regions and enhancer hotspots associated with NASH prone (NP) livers. (A) Schematic for defining enhancer hotspots in NP livers. NP-specific enhancers were first stitched based on the distance criteria (< 12.5 kb). Stitched enhancers are then ranked based on the transcription activity (if single enhancer regions) or the sum of transcription activity from each constituent NP-specific enhancer (if multi-enhancer regions). (B) Sankey diagram showing that NP-specific enhancers form 42 multi-enhancer regions and 31 enhancer hotspots (EHs). (C) Ranked plot defining NP-specific EHs (n = 31) and non-EHs (n = 333). (D) Transcription fold change of genes associated with either NP-specific multi-enhancer or single enhancer regions. * P < 0.05 by unpaired two-sample Wilcoxon test. (E) Transcription fold change of genes associated with either NP-specific EHs or non-EHs. * P < 0.05 by unpaired two-sample Wilcoxon test. (F) NP-activated genes that are associated with NP-specific multi-enhancer regions and/or EHs are shown. Lollipop plot height indicates the distance between NP-specific multi-enhancer regions/EHs and the associated transcription start sites (TSSs). Dot size denotes the activity level (based on ChRO-seq signal) of NP-specific multi-enhancer regions/EHs. Also see Supplementary Table 5. (G-J) Transcription activity (normalized ChRO-seq signal), all TREs, and NP-specific EHs around the loci of *ATF3* (G) and *HES1* (H), MIR23AHG (primary transcript of miR-24-2, miR-27a and miR-23a) (I), and *CSRNP1* (J) are shown. NASH-prone (NP), n = 8.

Among all 364 stitched enhancers in NP, we identified 42 that contain multiple NP-specific enhancers (“NP-specific multi-enhancer regions”, **Figure 3B**). Of the 364, 31 were defined as NP-specific enhancer hotpots (**Figure 3B-C**). We assigned NP-specific multi-enhancer regions and enhancer hotspots to the closest annotated TSSs. The genes associated with either NP-specific multi-enhancer regions or enhancer hotspots are on average more transcriptionally active in NP livers than those associated with NP-specific single-enhancer regions or non-enhancer hotspots (**Figure 3D-E**). The NP-activated genes that are associated with NP-specific multi-enhancer regions or enhancer hotspots are shown in **Figure 3F** (n = 32). Among these genes are those encoding transcription factors ATF3(26, 54) (**Figure 3G**), AP-1 (*FOS*, *JUNB*)(55) and the Notch signaling effector HES1(56, 57) (**Figure 3H**), all of which have been reported as key molecular players in hepatic fibrosis in humans or *in vivo* mouse models. Also, although activation of Wnt signaling is known to drive liver fibrosis(45), the downstream key mediators are not completely understood. In this analysis we have uncovered at least two key Wnt signaling mediators, *EXT1*(58) and *CSRNP1*(59, 60) (**Figure 3J**), neither of which has been previously linked to NAFLD or NASH and thus warrant detailed investigation. In a separate analysis, we also assigned NP-specific multi-enhancer regions or enhancer hotspots to the closest putative TSSs for miRNAs (methods). We found that miR-27a/miR-24-2/miR-23a, which has been linked previously to dysregulation of hepatic lipid metabolism and viral hepatitis in humans(61–63), is the only microRNA locus that is associated with NP-specific enhancer hotspots in our analysis (**Figure 3I**). Taken together, we have identified a small subset of NP-activated genes that are associated with NP-specific multi-enhancer regions or enhancer hotspots, which represents those genes with potentially critical etiological roles in NASH development.

Similar analyses were performed with NR-specific enhancers. Among all 608 stitched enhancers in NR, we identified 77 that contain multiple NR-specific enhancers (“NR-specific multi-enhancer regions”, **Figure 4A**). Of the 608, 57 were defined as NR-specific enhancer hotspots (**Figure 4A-B**). The genes associated with either NR-specific multi-enhancer regions or enhancer hotspots are on average more transcriptionally active in NR livers than those associated with NR-specific single-enhancer regions or non-enhancer hotspots (**Figure 4C-D**). The NR-activated genes that are associated with NR-specific multi-enhancer regions or enhancer hotspots are shown in **Figure 4E** (n = 31). These include *SESN2*(*64*) (ER stress regulator) and *PRKAB2* (AMPK complex subunit; **Figure 4F**)(65), both of which have been recently demonstrated to prevent liver damage and/or NASH progression. This analysis also highlighted several hepatocyte metabolic genes (e.g., *HAO2*, *ACAA2*, *GYS2*; **Figure 4E, G**), which is consistent with a recent observation that liver fibrosis is accompanied by loss of hepatocyte cellular identify and function(25). Furthermore, our analysis uncovered genes (notably *GATM* and *GLYAT*) which encode proteins that utilize glycine to carry out various biological processes. GATM, which mediates creatine biosynthesis from glycine, and GLYAT (**Figure 4H**), which conjugates glycine with acyl-CoA species during xenobiotic and lipid metabolism(66), could be critical in contributing to NASH resistance in humans.

**Figure 4.**
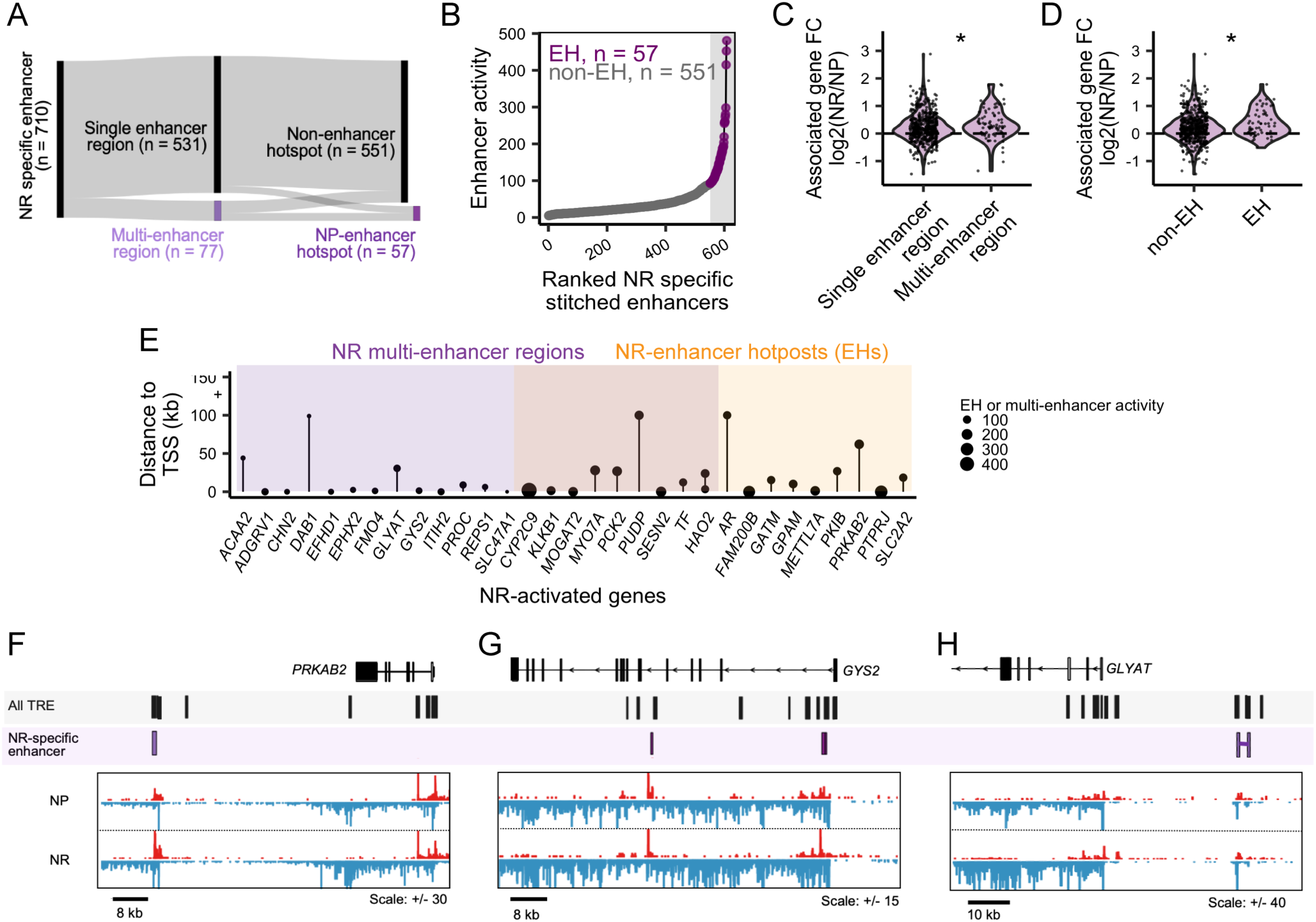
Identification of multi-enhancer regions and enhancer hotspots associated with NASH resistant (NR) livers. (A) Sankey diagram showing that NR-specific enhancers form 77 multi-enhancer regions and 57 enhancer hotspots (EHs). (B) Ranked plot defining NR-specific EHs (n = 57) and non-EHs (n = 551). (C) Transcription fold change of genes associated with either NR-specific multi-enhancer or single enhancer regions. * P < 0.05 by unpaired two-sample Wilcoxon test. (D) Transcription fold change of genes associated with either NR-specific EHs or non-EHs. * P < 0.05 by unpaired two-sample Wilcoxon test. (E) NR-activated genes that are associated with NR-specific multi-enhancer regions and/or EHs are shown. Lollipop plot height indicates the distance between NR- specific multi-enhancer regions/EHs and the associated transcription start sites (TSSs). Dot size denotes the activity of NR-specific multi-enhancer regions/EHs. Also see Supplementary Table 5. (F-H) Transcription activity (normalized ChRO-seq signal), all TREs, and NR-specific EHs around the loci of *PRKAB2* (F), *GYS2* (G), and *GLYAT* (H) are shown. NASH-resistant (NR), n = 6.

### Active TF gene networks in NP livers

TFs initiate transcription of downstream genes by binding to sequence-specific motifs present within nearby active enhancers. To identify candidate master TF regulators relevant to NP livers, we performed motif enrichment analysis within NP-specific enhancers (**Figure 5A**). The TFs that are highly expressed in NP livers (normalized counts > 500 in NP) and exhibit significantly enriched binding motifs in NP-specific enhancers (p < 0.05, q-value < 0.05, enrichment fold change > 1.5, target sequences with motif > 10% by HOMER) are shown in **Supplementary Figure 2**. The NP associated TFs include ATF members (e.g., ATF3)(26, 67), AP-1 factors (e.g., JUN, JUNB, FOSL2)(55, 68), STAT members (e.g., STAT3)(69, 70), CEBPB(71), and ELF3(25) (**Figure 5B**), several of which have been implicated in mouse models of NAFLD formation and/or disease progression to NASH. Notably, we observed that many of the NP-associated TFs are themselves significantly activated at the transcriptional level in NP compared to NR (**Figure 5C**), and even associated with NP-specific enhancer hotspots (e.g., AP-1 factors, ATF3), which suggests that activation of TF networks in NP livers is achieved by transcriptional activation of NP associated TFs as well as the enhancers containing the corresponding binding motifs.

**Figure 5.**
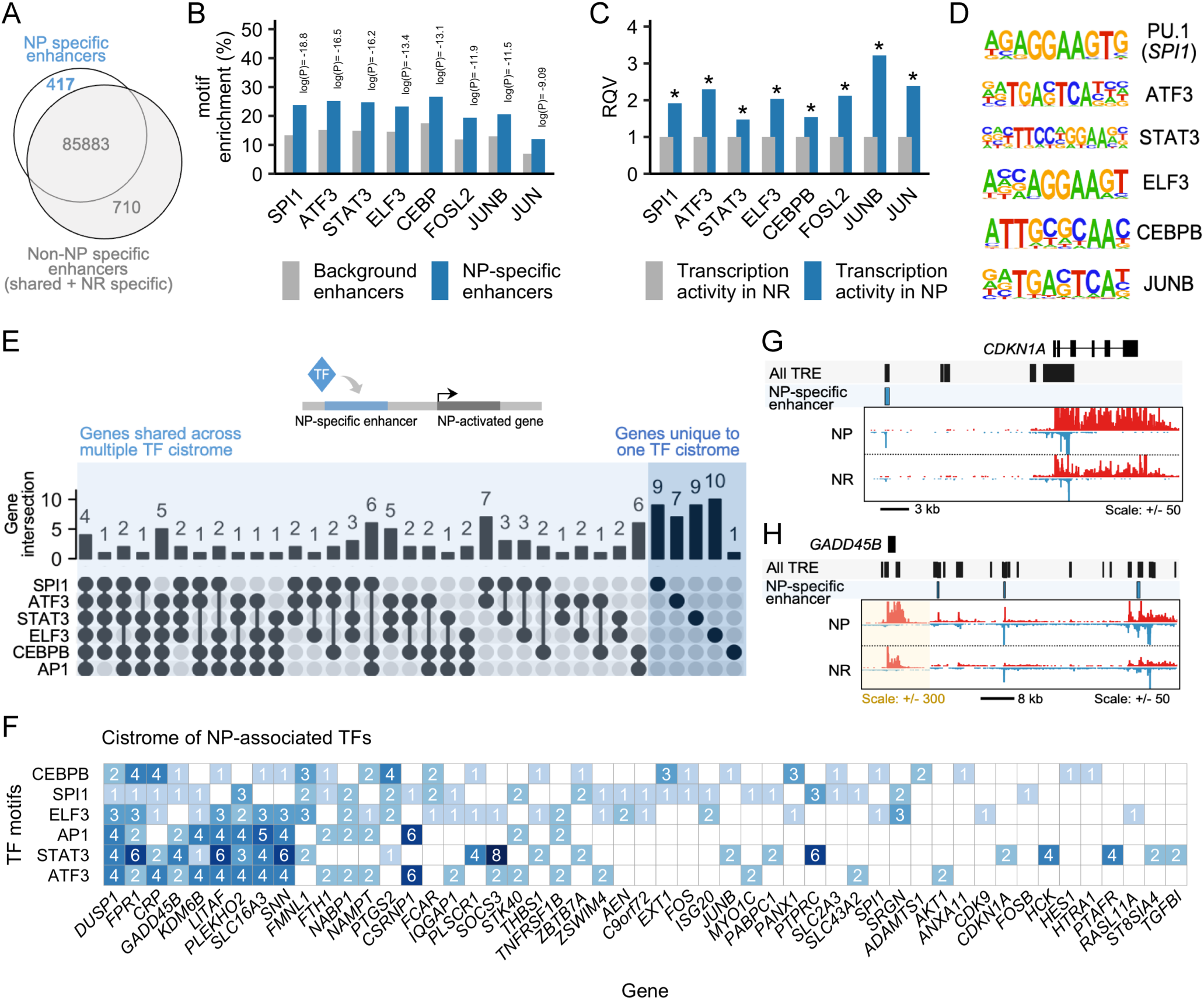
Key transcription factor (TF) gene networks in NASH prone (NP) livers. (A) Venn diagram indicating selection of NP-specific enhancers for motif enrichment analysis. (B) TF motif enrichment in NP-specific enhancers (p < 0.05, q < 0.05, enrichment foldchange > 1.5, target sequences with motif > 10% by HOMER). (C) Transcription fold-change of TFs highlighted in (B). * = P < 0.05, adjusted P < 0.2, log2 foldchange < 0 in NR compared to NP (Wald test; DESeq2). (D) HOMER motifs of NP-associated TFs. AP-1 motifs include JUNB, JUN and FOSL2. JUNB is used as a representative motif in this figure panel. (E) Upset plot showing NP-activated genes associated with one or more NP-associated TFs: SPI.1, ATF3, STAT3, ELF3, AP-1 (JUNB, JUN, or FOSL2) and CEBPB. (F) NP-activated genes associated with one or more NP-associated TFs. Motif counts are shown. See the complete list in Supplementary Table 6. (G-H) Transcription activity (normalized ChRO-seq signal), all TREs, and NP-specific enhancers around loci of *CDKN1A* (F) and *GADD45B* (G) are shown. NASH-prone (NP), n = 8.

We next sought to define major TF-gene networks (cistromes) in NP livers. Specifically, we selected SPI1, ATF3, AP-1 (including FOS2L, JUNB, and JUN), STAT3, CEBPB and ELF3 as TFs of interest due in part to their significant activation at the transcriptional level (**Figure 5C**). We then identified NP-activated genes that are associated with NP-specific enhancers (within +/-100 kb window from TSS) containing one or more of the aforementioned binding motifs (**Figure 5D-E; Supplementary Table 6**). Through this analysis, we identified genes that are shared across different TF cistromes or uniquely associated with a particular TF (**Figure 5E-F**). For example, *HES1*, an NP-specific enhancer hotspot linked gene (**Figure 3H**), is uniquely associated with the CEBPB motif (**Figure 5E-F**). Another NP-specific enhancer hotspot linked gene, *CSRNP1* (**Figure 3J**), is shared across the networks of ATF3, AP-1, and SPI1 (**Figure 5E-F**). In addition, we found that the *SOCS3* locus is associated with NP-specific enhancers that harbor a very large number of STAT3 motifs (**Figure 5F**). This is consistent with previous observations of *SOCS3* as a direct target of STAT3 and the co-activation of STAT3 and SOCS3 in a human liver organoid model of fibrosis(72). We noticed that several genes in Figure 5F are direct targets of p53, which plays an essential role in driving apoptosis in NASH(47, 73, 74). Our analysis revealed that *CDKN1A* (p21) is uniquely associated with the STAT3 cistrome (**Figure 5F, G**), whereas other p53 targets (e.g., *THBS1*, *AEN*, and *GADD45B*) are associated with multiple TF cistromes (**Figure 5F, H**).

### Active TF gene networks in NR livers

To identify candidate master TF regulators relevant to NR livers, we performed motif enrichment analysis within NR-specific enhancers (**Figure 6A**). The analysis revealed only four TFs that are highly expressed in NR livers (normalized counts > 500 in NR) and exhibit significantly enriched binding motifs in NR-specific enhancers (p < 0.05, q-value < 0.05, enrichment fold change > 1.5, target sequences with motifs > 10% by HOMER) are defined as NR associated TFs (**Figure 6B**). The 4 identified NR associated TFs, namely HNF4A(47), HNF1B(75), FOXA2(76) and PPARA(77, 78), have reported roles in preventing NAFLD/NASH development in various rodent models. Interestingly, the genes encoding all of the NR associated TFs are transcribed at similar levels between NR and NP (**Figure 6C**). This suggests that activation of NR-associated TF networks relies primarily on enhancing the activity of the TFs at enhancers without affecting their own transcription levels, unlike what was observed in NP (**Figure 5C**).

**Figure 6.**
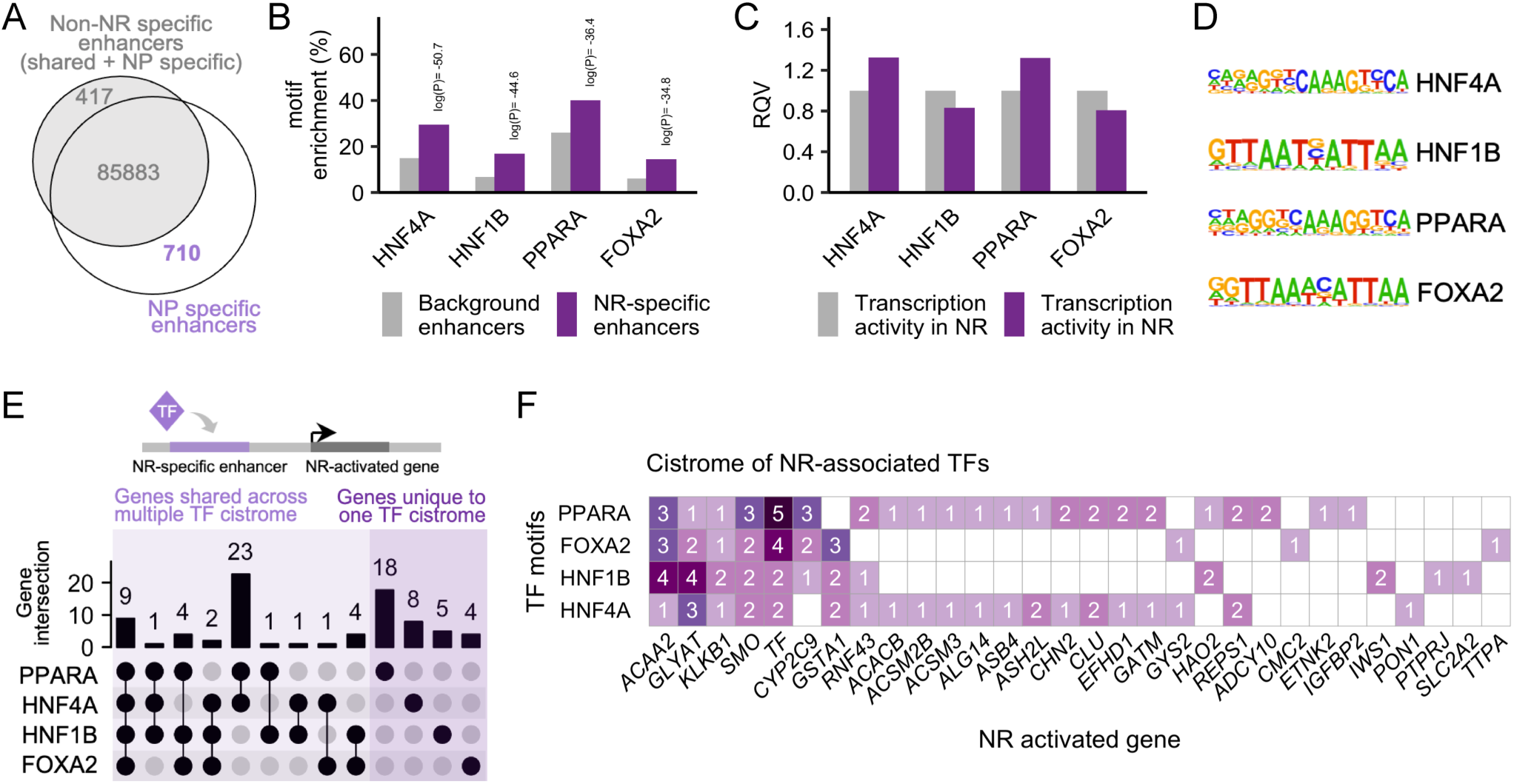
Key transcription factor (TF) gene networks in NASH resistant (NR) livers. (A) Venn diagram indicating selection of NR-specific enhancers for motif enrichment analysis. (B) TF motif enrichment in NR-specific enhancers (p < 0.05, q < 0.05, enrichment foldchange > 1.5, target sequences with motif > 10% by HOMER). (C) Transcription fold-change of TFs highlighted in (B). (D) HOMER motifs of NR-associated TFs. (E) Upset plot showing NR-activated genes associated with one or more NR-associated TFs: HNF4A, HNF1B, PPARA, and FOXA2. (F) NR-activated genes associated with one or more NR-associated TFs. Motif counts are shown. See the complete list in Supplementary Table 6. NASH-resistant (NR), n = 6.

Next, to define TF-gene networks (cistrome) in NR livers, we identified NR-activated genes that are associated with NR-specific enhancers containing one or more of the aforementioned binding motifs (**Figure 6D-E; Supplementary Table 6**). This analysis identified genes that were known direct targets of the TFs of interest, such *IGFBP2* (PPARA target)(*79*) and *GYS2* (HNF4A target)(*80*) (**Figure 6F**). Moreover, we uncovered previously undescribed TF-gene interactions potentially critical for NASH resistance that warrant future validation. For example, we showed that *GLYAT* and *TF* (**Figure 5E,H**) are shared across all four NR-associated TFs (**Figure 6F**), whereas *GATM* (**Figure 5E**) is likely under the direct control of only HNF4A and PPARA (**Figure 6F**).

### Intersection analysis of disease/trait-SNPs and NP/NR-specific enhancers

Genome-wide association studies (GWAS) have uncovered thousands of genetic variants associated with complex disease traits; however, most of them are present in non-coding regions with unclear roles in disease etiology. To explore potential single nucleotide polymorphism (SNP)-containing enhancers that may be relevant to NP or NR phenotype, we intersected NP- and NR-specific active enhancer regions with the locations of tag SNPs (and those in linkage disequilibrium) that are associated with NAFLD and related phenotypic traits (GWAS catalog**; Supplementary Table 7**). The analysis showed that *rs1205* (known associated trait: C-reactive protein level) is present in a NP-specific enhancer region (**Figure 7A**). We also found that *rs3847302* (known associated trait: HDL cholesterol) is located within a NR-specific enhancer region (**Figure 7B**). We also performed follow-up TF motif analysis (using MEME suite) to identify candidate TFs that could have disrupted binding due to different allelic versions of SNPs within these enhancers (**Supplementary Table 8**). The analysis revealed that *rs3847302* may interrupt enhancer binding with several TFs, most notably HNF4A (q-value = 0.0107 by MEME suite; which we defined as one of the four top NR-associated TFs).

**Figure 7.**
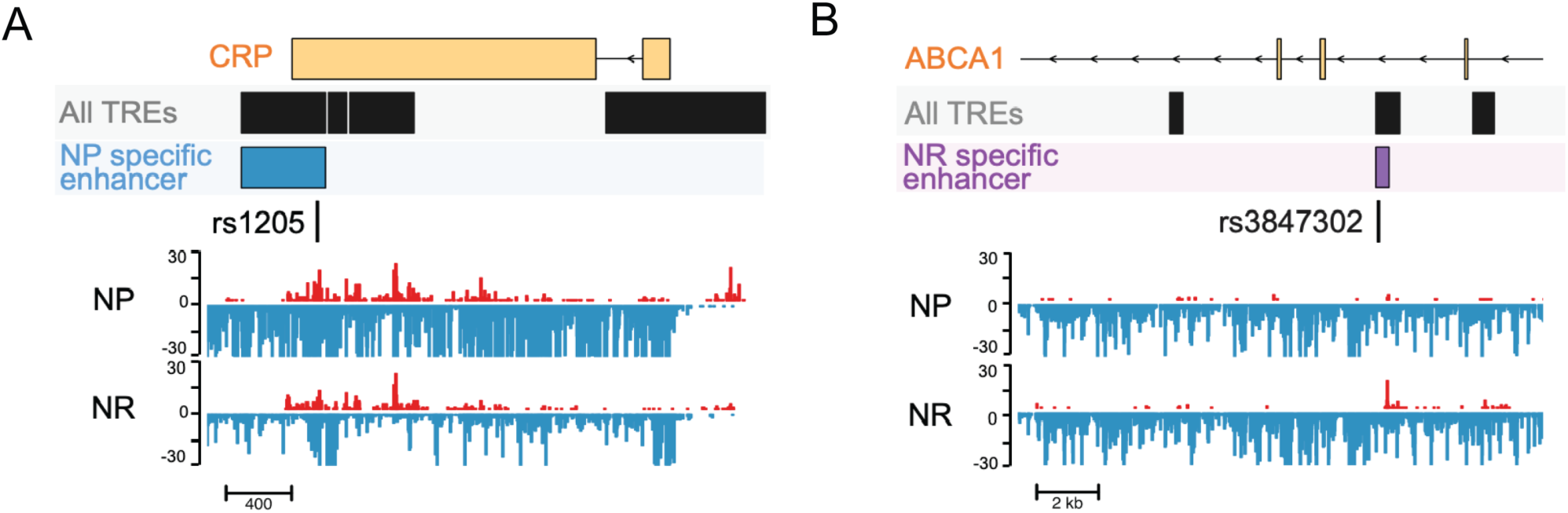
Intersection analysis of disease/trait-SNPs and NP/NR-specific enhancers. (A) Genomic location and transcriptional activity of NP-specific enhancer containing *rs1205* (chr1:159711980-159712490; hg38). (B) Genomic location and transcriptional activity of NR-specific enhancer containing *rs3847302* (chr9:104886231-104886650; hg38). NASH-resistant (NR), n = 6; NASH-prone (NP), n = 8.

### NASH resistance is associated with elevated circulating serine and activation of hepatic serine synthesis and glycine utilization pathways

The identification of genes that code for glycine utilization factors (GATM and GLYAT) across multiple analyses of NR-specific enhancers led us to focus next on amino acid metabolism. Given that circulating amino acid profiles can be influenced by the status of hepatic amino acid metabolism(66, 81, 82), we measured amino acids and related metabolites in the blood of NR and NP subjects (**Figure 8A**). We observed that NR subjects exhibit a trend toward higher circulating glycine (27% increase; p = 0.14) and a significant increase in circulating serine (38% increase; p = 0.028) compared to NP subjects (**Figure 8A**). Moreover, we showed that circulating glycine is positively correlated with circulating serine but not the other amino acids across all the samples in this study (**Figure 8B**), suggesting a strong relationship between glycine and serine metabolism in the context of NASH development.

**Figure 8.**
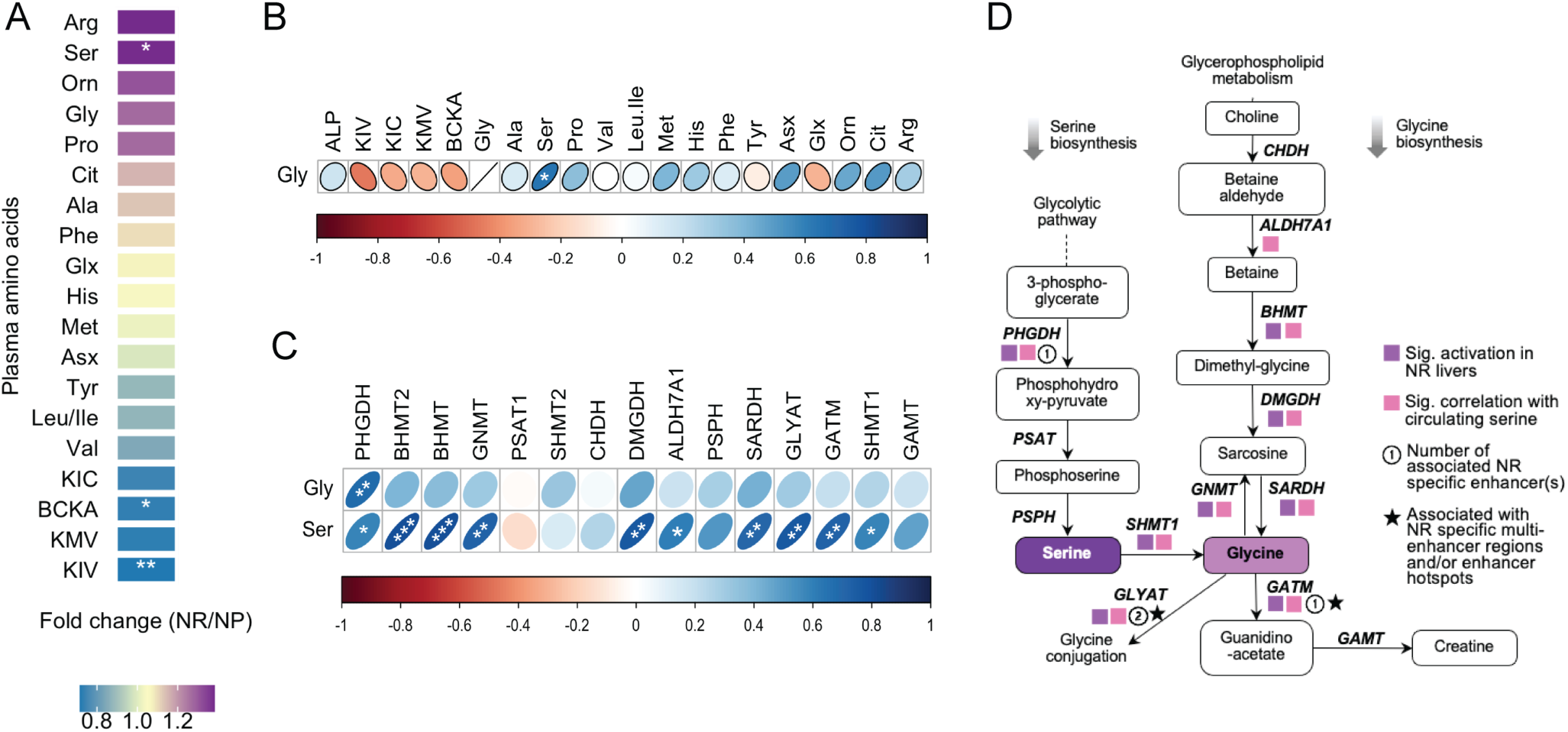
Human NR livers are marked by a robust transcriptional activation of genes in serine and glycine biosynthesis and utilization. (A) Circulating amino acid levels of NR compared with NP subjects. Color shade indicates fold change in NR relative to NP subjects. * P < 0.05, ** P < 0.05. (B) Pearson correlation heatmap of circulating glycine (Gly) with all the other circulating amino acids. (C) Pearson correlation heatmap of circulating Gly and serine (Ser) with hepatic ChRO-seq levels of genes involved in serine/glycine biosynthesis and utilization. In (B-C), color shade denotes correlation coefficient. * P < 0.05, ** P < 0.01 and *** P < 0.001. (D) Summary of genes involved in serine/glycine biosynthesis and utilization in human NP and NR livers. NASH-resistant (NR), n = 6; NASH-prone (NP), n = 8.

Serine is known as an important substrate for glycine biosynthesis. The increase in circulating serine in NR subjects (**Figure 8A**) motivated us to examine further the pathway of glycine, serine and threonine (GST) synthesis and metabolism (KEGG pathway hsa00260). Our analyses led to several interesting observations. First, circulating serine circulating significantly correlated with many genes in GST metabolism, while circulating glycine significantly correlated with only *PHGDH* across all the liver samples in our study (**Figure 8C-D**). Second, compared to NP livers, NR livers exhibit robust transcriptional activation of genes in pathways related to serine biosynthesis (e.g., *PHGDH*), serine-glycine conversion (e.g., *SHMT1*), glycine synthesis (*BHMT*(83), *DMGHD* and *SARDH*) (**Figure 8D**). As shown in our previous analysis, genes involved in glycine utilization (GLYAT, GATM) were significantly activated in NR livers (**Figure 8D**) and associated with NR-specific multi-enhancer regions (**Figure 4**). Therefore, the robust production of serine and glycine likely supports glycine utilization in NR livers (summarized in **Figure 8D**). Overall, these findings indicate that circulating serine could serve as an indicator of hepatic glycine metabolism, and that enhanced glycine utilization may represent a critical mechanism underlying NASH resistance.

## Discussion

In this study, ChRO-seq was leveraged to define for the first time the active enhancer signature of NASH-prone (NP) and NASH-resistant (NR) phenotypes in humans with severe obesity. Although whole-transcriptome profiling has been used previously to characterize the gene expression landscape of NAFLD/NASH in humans(14, 15, 19), the chromatin activity patterns, enhancer signature, and TF networks that drive gene expression in NASH remain largely undefined. The results of this study fill this important knowledge gap. In addition, while numerous studies in animal models have reported a wide array of genes that may confer beneficial effects in the liver toward mitigating NASH, which of them (if any) are relevant to humans has remained unclear. An important contribution of our study is to help determine which of these many candidates are most pertinent in humans.

The present study highlighted genes that are candidate key drivers of the NP phenotype. These genes include pro-apoptotic factors such as *CDKN1A* and *GADD45B*, the former of which has been linked to NAFLD progression(84) but the latter of which has not previously been reported in the context of NAFLD/NASH and therefore warrants detailed characterization. Our analyses also provided new insights into regulatory mechanisms of genes that have been linked to NAFLD progression in human patients. For example, we showed that *LITAF*, which is elevated in the livers of children with NASH(51), is shared across gene regulatory networks of multiple NP-associated TFs. In another example, we linked an NP-enhancer hotspot to tyrosine kinase *SYK*, the hepatic expression of which positively correlates with fibrosis stage in human tissues(85). NP-specific enhancer associated genes that were previously reported in cell/animal models but reported in our study for the first time in humans include: *ARID5B* (hepatic stellate cell activation(86)), *ARRB2* (fibrosis(87)), *FMNL1* (macrophage migration(88)), and *PANX1* (apoptosis and fibrosis(89–91)). We also uncovered novel genes that had not been previously linked to NASH progression in humans or animal models. One example is *SNN*, the locus of which is associated with NP-specific enhancers that harbor a large number of motifs of multiple NP-associated TFs, pointing to a potentially important role in NASH development. SNN is known as an apoptosis mediator and linked to trimethyltin toxicity (92); however, its role in NAFLD/NASH pathogenesis remains unexplored and warrants future characterization.

One of the findings in this study is the identity of the TFs that are most strongly associated with NP, some of which have been shown to play a critical role in NASH development in various rodent models. Mice with genetic ablation of CEBPB exhibit significant reductions of hepatic lipid accumulation and inflammation compared to wild type mice in response to a methionine and choline deficient diet (MCD)(71). More recently, ELF3, JUN and FOSL2 were discovered as part of the fibrosis-activated TF network that drives transcriptional rewiring in hepatocytes of mice fed two different NASH-inducing diets(25). In addition, the current literature indicates that some of the NP-associated TFs uncovered in our study exert cell-type specific roles in the liver during NASH progression. For example, in mouse NAFLD/NASH models, Jun/AP-1 was observed to promote cell survival in hepatocytes but drive pro-inflammatory/pro-fibrotic programs in non-parenchymal cells(24, 55). Another notable example is HES1, which has been shown to exhibit cell type specific expression patterns in human livers (hepatocytes vs. non-parenchymal cells) over the course of disease progression from healthy to steatosis to NASH(56). Active enhancer profiles, which are known to be cell-type specific, may be a key underlying mechanism for directing cell-type specific roles of TFs. In the future, it will be critical to extend the work of this study to achieve single-cell resolution on TF networks.

Our study also identified genes that may be pivotal for NASH resistance in humans with obesity and defined HNF4A, HNF1B, FOXA2 and PPARA as the core NR-associated TFs. While these TFs are already known to be important for several liver functions, we pinpoint in this study specific downstream genes that are likely especially critical for driving the NR phenotype in humans. For example, *PON1*(93), *ETNK2*(15) and *IGFBP2*(94) have been reported as promising markers that distinguish NASH from simple steatosis or healthy livers. Our analysis also uncovered which among the genes previously implicated in mouse models of NAFLD/NASH are likely most relevant in humans. For example, a mouse study described the critical role of *Smo* in preventing injury-induced apoptosis(95). We showed that *SMO* is transcriptionally activated in NR livers, associated with NR-specific enhancers, and shared across gene networks of all the NR-associated TFs defined in our study. In addition, liver-specific overexpression of *Clu* in mice was shown to significantly attenuate MCD-induced NASH development(96). Consistent with this observation, we identified *CLU* as an NR-specific enhancer associated gene most likely under the control of HNF4A and PPARA. Most importantly, our study also revealed novel genes that are associated with NR-specific enhancers and may play critical roles in contributing to NASH resistance. An example is *TTPA*, the genetic ablation of which was reported to enhance inflammatory responses in mice, though not in the context of NAFLD/NASH(97). Another example is *PTPRJ* (coding CD148), the deficiency of which has been shown to promote pulmonary fibrosis(98). Further work is required to validate the NR-specific super enhancers and TF networks identified here and investigate their roles in NAFLD/NASH pathogenesis.

The most intriguing biological processes that we identified to be strongly associated with NASH resistance are glycine conjugation-mediated acyl-CoA clearance (catalyzed by GLYAT) and glycine-based creatine synthesis (catalyzed by GATM). GLYAT is associated with multiple NR-specific enhancers and GATM is associated with an NR-specific enhancer hotspot. Both enzymes require glycine as a substrate to carry out their functions, pointing to the importance of glycine supply in NR livers. Indeed, our data showed that genes involved in glycine synthesis (e.g., *BHMT*, *SARDH*), serine synthesis (e.g., *PHGDH*) as well as serine conversion to glycine (e.g., *SHMT*) are all significantly activated in NR livers, many of which are positively correlated with circulating serine levels. Our findings are consistent with two human studies, one of which indicated serine deficiency in human NAFLD/NASH patients(99) and the other of which showed that glycine deficiency is a risk factor for liver fibrosis(100). Our observations also support recent studies that suggest serine- or glycine-based treatments for NASH prevention(101, 102). The novelty of the present study is that we pinpointed particularly important genes and biological processes that mediate the beneficial effects of serine and glycine in the liver.

It is important to highlight the added value of our first-ever super enhancer analysis of NP and NR livers. Studies that perform gene expression analysis (e.g., RNA-seq) will identify hundreds to thousands of genes that are differentially expressed between NP and NR. However, in such studies, it is often unclear which genes are the most critical for driving the phenotypic differences. By leveraging the observation that super enhancers typically mark genes with particularly critical roles in defining cellular behavior(22, 31, 32), we are able to define a small number of genes (n=32 in **Figure 3F** and n=31 in **Figure 4E**), and their likely control elements, that merit detailed functional investigation, which in turn may guide future therapeutic development for NASH.

Overall, by leveraging the cutting-edge genome-scale technique ChRO-seq on human liver biopsies from a cohort of individuals with obesity, we defined for the first-time active enhancer signatures that distinguish the NR from the NP phenotype. We were able to identify genes that are linked to NP- or NR-specific enhancers/enhancer hotspots and offer a basis for understanding how enhancer status is modulated to regulate transcriptional activity of genes during NAFLD progression. We also describe the re-wiring of TF networks in NASH relative to NAFLD. Some of these findings in this study help uncover the specific genes and networks reported in animal models that may be most relevant in humans. Other findings reveal novel candidate molecular factors that have not previously been linked to NAFLD/NASH and may serve as potential therapeutic targets for managing NASH.

## Methods

### Human sample description

Tissue specimens, blood, and plasma samples were obtained from the Biobank of the Institut Universitaire de Cardiologie et de Pneumologie de Québec (Québec Heart and Lung Institute, QHLI, Québec City, QC, Canada) according to institutionally approved management modalities. Participants were recruited during a preoperative visit for bariatric surgery and met the inclusion/exclusion criteria for the procedure (103). The study population included 18 female patients of European ancestry with severe obesity (BMI > 45 kg/m^2^) aged 29-56 years from the eastern provinces of Canada who underwent bariatric surgery at the Institut Universitaire de Cardiologie et de Pneumologie de Québec (Québec Heart and Lung Institute, QHLI, Québec City, QC, Canada) and were not on diabetes medications. Blood samples were obtained at the time of admission for surgery. Liver tissues were obtained during bariatric surgery by incisional biopsy of the left lobe and were not cauterized. The sampling procedure and position was standardized among surgeons. A portion of the liver tissues were rapidly frozen in liquid nitrogen and another portion was fixed for histology grading by trained pathologists according to (104). The participants used for the study were selected based on the presence or absence of NASH defined as steatosis alongside both lobular inflammation and ballooning. 6 out of the 18 samples showed no indication of hepatocyte ballooning and hepatic fibrosis were categorized as NASH resistant (NR). 8 out of the 18 samples with indications of hepatocyte ballooning, hepatic fibrosis, lobular inflammation, and portal inflammation were categorized as NASH prone (NP). The remaining 4 samples in this cohort satisfied neither NR nor NP criteria and thus were excluded from NR vs. NP comparison throughout the entire study. Subjects in NR and NP groups were matched for BMI, sex, glucose tolerance, PNPLA3 genotype and smoking status. Additional sample and subject details are provided in Supplementary Table 1.

### Genotyping

Patatin-like phospholipase domain-containing protein 3 (PNPLA3) genotyping for the Ile148Met variant associated with hepatic steatosis (rs738409) was performed on genomic DNA extracted from the blood buffy coat using the GenElute Blood Genomic DNA kit (Sigma, St. Louis, MO, USA). rs738409 was genotyped using validated primers and TaqMan probes (Applied Biosystems, Waltham, MA). PNPLA3 genotypes were determined using 7500 Fast Real-Time PCR System (Applied Biosystems, Waltham, MA).

### Plasma metabolite measurements

Plasma amino acids were measured by targeted metabolomics methods, as previously described(105, 106). Briefly, plasma amino acid profiling was performed by tandem mass spectrometry (MS/MS). All MS analyses employed stable-isotope-dilution with internal standards from Isotec, Cambridge Isotopes Laboratories, and CDN Isotopes.

### Chromatin isolation

Chromatin isolation for length extension chromatin run-on sequencing (ChRO-seq) was performed as previously described(28, 31). To isolate chromatin from pulverized frozen liver biopsies, samples were incubated in 1X NUN buffer [20 mM HEPES, 7.5 mM MgCl2, 0.2 mM EDTA, 0.3 M NaCl, 1M urea, 1% NP-40, 1 mM DTT, 50 units/mL RNase Cocktail Enzyme Mix (Thermo Fisher Scientific, Waltham, MA; AM2286), 1X Protease Inhibitor Cocktail (Roche, Basel, Switzerland; 11873580001)] in an Eppendorf Thermomixer (Eppendorf, Hamburg, Germany) set at 12°C and 2000 rpm for 30 minutes. The chromatin was pelleted by centrifugation at 12,500 × g for 30 minutes at 4°C, washed by 50 mM Tris-HCl (pH 7.5) containing 40 units/mL SUPERase inhibitor and stored in chromatin storage buffer (50 mM Tris-HCl pH 8.0, 25% glycerol, 5 mM magnesium acetate, 0.1 mM EDTA, 5 mM DTT, and 40 units/mL SUPERase In RNase Inhibitor). To solubilize the chromatin into the storage buffer, samples were loaded into a Bioruptor (Diagenode, Denville, NJ) and sonicated with repeated cycles (10 minutes per cycle, consisted of 10 rounds of 30s on and 30s off). Chromatin solubilized in storage buffer was stored at −80°C until usage for library construction.

### ChRO-seq library preparation

Library preparation for length extension ChRO-seq was performed as previously described(28, 31). To perform run-on reaction with the solubilized chromatin, samples were mixed with an equal volume of 2X run-on reaction mix [10 mM Tris-HCl pH 8.0, 5 mM MgCl2, 1 mM DTT, 300 mM KCl, 400 μM ATP, 0.8 μM CTP, 400 μM GTP, 400 μM UTP, 40 μM Biotin-11-CTP (Perkin Elmer, Waltham, MA; NEL542001EA), 100 ng yeast tRNA (VWR, 80054–306), 0.8 units/μL SUPERase In RNase Inhibitor, 1% (w/v) Sarkosyl]. The run-on reaction was incubated in an Eppendorf Thermomixer at 37°C for 5 min (700 rpm) and stopped by adding Trizol LS (Life Technologies, Carlsbad, CA; 10296–010). RNA samples were precipitated by GlycoBlue (Ambion, Austin, TX; AM9515) and resuspended in diethylpyrocarbonate (DEPC)-treated water. To perform base hydrolysis reaction, RNA samples in DEPC water were heat denatured at 65°C for 40 s and incubated in 0.2N NaOH on ice for 4 min. Base hydrolysis reaction was stopped by neutralizing with Tris-HCl pH 6.8. Nascent RNA was enriched using streptavidin beads (NEB, Ipswich, MA; S1421) followed by RNA extraction using Trizol. To perform adapter ligations, nascent RNA samples were processed through the following steps: (i) 3′ adapter ligation with T4 RNA Ligase 1 (NEB, Ipswich, MA; M0204), (ii) RNA binding with streptavidin beads (NEB, Ipswich, MA; S1421) followed by RNA extraction with Trizol, (iii) 5′ de-capping with RNA 5′ pyrophosphohydrolase (NEB, Ipswich, MA; M0356), (iv) 5′ end phosphorylation using T4 polynucleotide kinase (NEB, Ipswich, MA; M0201), (iv) 5′ adapter ligation with T4 RNA Ligase 1 (NEB, Ipswich, MA; M0204). The 5’ adaptor contained a 6-nucleotide unique molecular identifier (UMI) to allow for bioinformatic detection and elimination of PCR duplicates. Streptavidin bead binding followed by Trizol RNA extraction was performed again before final library construction. To generate ChRO-seq library, cDNA was generated through a reverse-transcription reaction using Superscript III Reverse Transcriptase (Life Technologies, Carlsbad, CA; 18080–044) and amplified using Q5 High-Fidelity DNA Polymerase (NEB, Ipswich, MA; M0491). Finally, ChRO-seq libraries were sequenced (5′ single end; single-end 75×) using the NextSeq500 high-throughput sequencing system (Illumina, San Diego, CA) at the Cornell University Biotechnology Resource Center. Raw and processed ChRO-seq data was deposited to NCBI Gene Expression Omnibus.

### ChRO-seq mapping and visualization

ChRO-seq mapping was performed with an established pipeline(107). Briefly, read quality was assessed using FastQC. PCR deduplication was performed by collapsing UMIs using PRINSEQ lite 0.20.2(108). Adapters were trimmed from the 3’ end of remaining reads using cutadapt 1.16 with a maximum 10% error rate, minimum 2 bp overlap, and minimum 20 quality score. Processed reads with a minimum length of 15 bp were mapped to the hg38 genome modified with the addition of a single copy of the human Pol I ribosomal RNA complete repeating unit (GenBank: U13369.1) using Burrows-Wheeler Aligner (BWA) 0.7.13(109). The location of the RNA polymerase active site was represented by a single base that denotes the 5’ end of the nascent RNA, which corresponds to the position on the 3’ end of each sequenced read. Supplementary Table 2 provides the mapping statistics of the ChRO-seq experiments. As denoted in the table, 4 samples in this patient cohort did not satisfy the criteria for NR or NP category based on their clinical profiles. The ChRO-seq data of these 4 samples were used only in the dREG analysis for calling active promoters and enhancers (see method section: Transcription activity transcriptional regulatory elements), but not elsewhere. To visualize ChRO-seq signal, data were converted to bigwig format using bedtools and UCSC bedGraphToBigWig. Bigwig files within a sample category were then merged and normalized to a total signal of 1×10^6^ reads. Genomic loci snapshots were generated using R package Gviz(110).

### Transcription activity of genes

Quantification of gene transcription activity was based on hg38 GENCODE v25 annotations. Read counts of gene loci were quantified using the R package bigwig (https://github.com/andrelmartins/bigWig). To quantify transcription activity of gene loci, stranded ChRO-seq reads within gene coordinates were counted, with exclusion of reads within 500 b downstream of transcription start site (TSS) to avoid bias generated by the RNA polymerase pausing at the promoters. Genes with gen body smaller than 1 kb were excluded from all the gene body related analysis. Principal components analysis (PCA) of genes was performed using counts with rlog transformation. In differential transcription analysis, ChRO-seq raw counts of gene loci were analyzed through DESeq2 1.30.1(111), which models based on negative binomial distribution, to define genes that are significantly activated in NASH-resistant or NASH-prone samples. Genes significantly activated in one specific group were defined by criteria of log2 foldchange > 0 (or < 0), normalized counts > 100, P < 0.05, padj < 0.2 (Wald test; DESeq2). Pathway analyses were performed using Enrichr(112–114) and KEGG pathway(115).

### Transcription activity transcriptional regulatory elements (TREs)

To identify active TREs across all the samples, bigwig files of the same strand from all samples were merged. This merged dataset was analyzed through dREG(107, 116), the peak-calling algorithm that detects short bidirectional expression patterns across the entire genome, to define active TREs. Classification of TREs into promoters and enhancers was based on hg38 GENCODE v25 annotations. TREs that had at least 1 b overlapping with the window of 1000 b upstream and 200 b downstream of TSS were defined as promoter regions; the remaining TREs were defined as enhancers. The R package bigwig (https://github.com/andrelmartins/bigWig) was used to quantify activity of TREs. To quantify the activity of TREs that reside entirely within annotated gene bodies, only reads on the opposite strand were used and multiplied by 2 in order to avoid the signal introduced by gene body transcription activity. To quantify the activity of TREs that do not reside entirely within annotated gene bodies, reads from both strands were used. PCA and hierarchical clustering analysis that computes pairwise correlations between samples based on promoter or enhancer profiles were performed using rlog transformation counts. To identify TREs that are uniquely active in NASH-resistant or NASH-prone samples, ChRO-seq raw counts of TREs were analyzed through DESeq2 1.30.1 (111). TREs associated with NASH-resistant phenotype were defined by criteria of log2 foldchange > 1 (or < −1), P < 0.05, padj < 0.2 (Wald test; DESeq2). TREs associated with NASH-prone phenotype were defined by criteria of log2 foldchange < −1), P < 0.05, padj < 0.2 (Wald test; DESeq2). In enhancer density analysis, the enhancers present within +/-100 kb window from annotated TSSs of genes (longest isoform if there are multiple; hg38 GENCODE v25) were counted.

### Transcription factor motif enrichment analysis

HOMER (117) was used to determine enrichment of sites corresponding to known motifs of a given set of enhancer peaks. Specifically, findMotifsGenome.pl function, with “given” as the size parameter, was used to query NASH-resistant enhancer peaks (using all enhancers that were not identified as NASH-resistant as background) and NASH-prone enhancer peaks (using all enhancers that were not identified as NASH-resistant as background). “Known motif” results generated from HOMER were reported in this study. To identify enhancers that contain binding motif(s) for a specific transcription factor, HOMER function annotatePeaks.pl (“given” as the size parameter; genome build hg38) was used.

### Defining enhancer hotspots

NASH-resistant and NASH-prone enhancer hotspots were defined using an analysis pipeline described previously(32). The enhancer hotspots in this study were identified by the criteria similar to the studies describing ‘super-enhancers’. Briefly, the enhancers uniquely activated in one specific group (NASH-resistant or -prone samples) were stitched based on proximity of distance (<12.5 kb). The stitched enhancers that contain multiple enhancers were define as ‘multi-enhancer regions’. To quantify the activity of stitched enhancers, counts of each enhancer were normalized by DESeq2 and summed for each stitched enhancer. To determine which stitched enhancers were qualified as ‘enhancer hotspots’, the stitched enhancers were first ranked by their activity, creating a curve of activity to rank plot. A tangent line was then applied to the curve. The ones above the cutoff point that is determined by the tangent line were defined as enhancer hotpots, whereas the ones below the cutoff point were non-enhancer hotspots. To identify candidate genes associated with phenotype-specific (NP or NR) multi-enhancer regions or enhancer hotspots, phenotype-specific multi-enhancer regions or enhancer hotspots were assigned to the closest annotated TSSs (GENCODE v25 annotations; only those genes with an average gene transcription activity > 100 normalized counts in the corresponding phenotype group were included). To identify candidate microRNA loci associated with phenotype-specific multi-enhancer regions or enhancer hotspots, the putative TSSs of microRNAs were curated from Sethupathy et al.(118)

### SNP intersection analysis

Using bedtools v2.29.2 intersect function, we intersected the locations of NR-(or NP-) specific enhancers with the locations of SNPs for traits relevant to liver functions or metabolic disorders. The R package gwascat (v2.18.0) was used to download disease/trait-associated SNPs from the NHGRI-EBI GWAS catalog(119). Variants associated with the following terms (p < 5 × 10^−8^) were extracted (ebicat38; gwascat): high-density lipoprotein cholesterol (HDLc), low-density lipoprotein cholesterol (LDLc), triglycerides (TG), cardiovascular disease, hypertension, type 2 diabetes (T2D), insulin, glucose, serum albumin, glycated hemoglobin (HbA1c), C-reactive protein (CRP), body mass index (BMI), bilirubin, liver enzymes, liver injury, and non-alcoholic fatty liver disease (NAFLD). The R package LDlinkR (v1.1.2) was used to query pairwise linkage disequilibrium (LD) between SNPs. R^2^ was calculated based on EUR populations (1000G project). The genomic coordinates of tag SNPs and SNPs in LD (R^2^ > 0.8) were converted from hg19 to hg38 using tool liftOver and used for the intersection analysis.

### Motif scan analysis

Motif identification with query sequences were performed using FIMO (Find individual motif occurrences)(^120^) in MEME suite 5.4.1(^121^). HOCOMOCO Human (v11 CORE) database was used as motif input.

### Statistics

Statistical analyses were performed using R (4.0.4). Statistical significance of blood measurements between groups was determined using unpaired two sample T-test. In sequencing studies, statistical significance was determined using DESeq2, where p values were calculated by Wald test and the p values were adjusted using Benjamin and Hochberg (BH) method. Other statistical methods including unpaired two-sample Wilcoxon test and two-sided Pearson correlation test were used as indicated in the figure legends. The p values in pathway enrichment analysis by Enrichr were calculated using Fisher’s exact test. Values of p < 0.05 (and adjusted p < 0.2 if in sequencing studies) were considered statistically significant unless otherwise noted. ∗ p < 0.05, ∗∗ p < 0.01, ∗∗∗ p < 0.001.

### Study approval

Tissue specimens, blood, and plasma samples were obtained from the Biobank of the Institut Universitaire de Cardiologie et de Pneumologie de Québec (Québec Heart and Lung Institute, QHLI, Québec City, QC, Canada) according to institutionally approved management modalities. All participants provided written, informed consent. All participants are identified by number and not by name or any protected health information (PHI).

## Acknowledgement

The authors acknowledge the invaluable collaboration of the surgery team, bariatric surgeons and Biobank staff of the IUCPQ. The authors also thank the Cornell University Biotechnology Resource Center for generating ChRO-seq genomic dataset.

## Author Contributions

Conceptualization: P.S., P.J.W., Y.-H.H.; Tissue collection and biobank resources: L.B., A.T.; Genotyping: M.C.V.; Metabolomics: O.I.; ChRO-seq library preparation: R.S.; Bioinformatic analyses and data curation: Y.-H.H.; Writing (original draft): Y.-H.H., P.S.; Review and editing: Y.-H.H., P.S., P.J.W., A.T.; Supervision: P.S., P.J.W.

