## Supplementary for "Super-enhancer signature reveals key mechanisms associated with resistance to non-alcoholic steatohepatitis in humans with obesity"

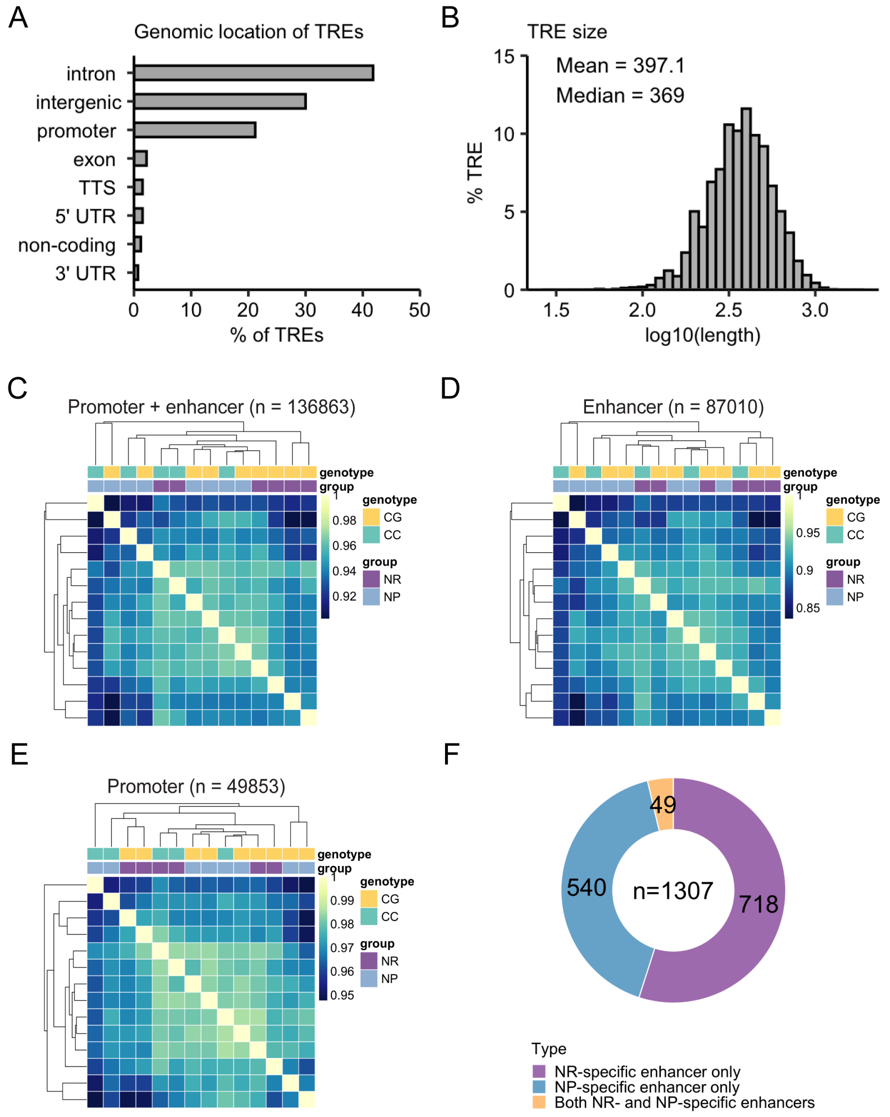


**Supplementary Figure 1.** (A) Genomic location of active transcriptional regulatory elements (TREs, n = 136863) identified across all samples by ChRO-seq. (B) Size distribution of TREs. (C) Hierarchical clustering analysis of all active TREs (enhancers and promoters) defined by ChRO-seq. (D) Hierarchical clustering analysis of active enhancers only. (E) Hierarchical clustering analysis of active promoters only. (F) Number of genes that are associated with both NR and NP enhancers, NR-specific enhancer(s) only, NP-specific enhancer(s) only. NASH-resistant (NR), n = 6; NASH-prone (NP), n = 8.

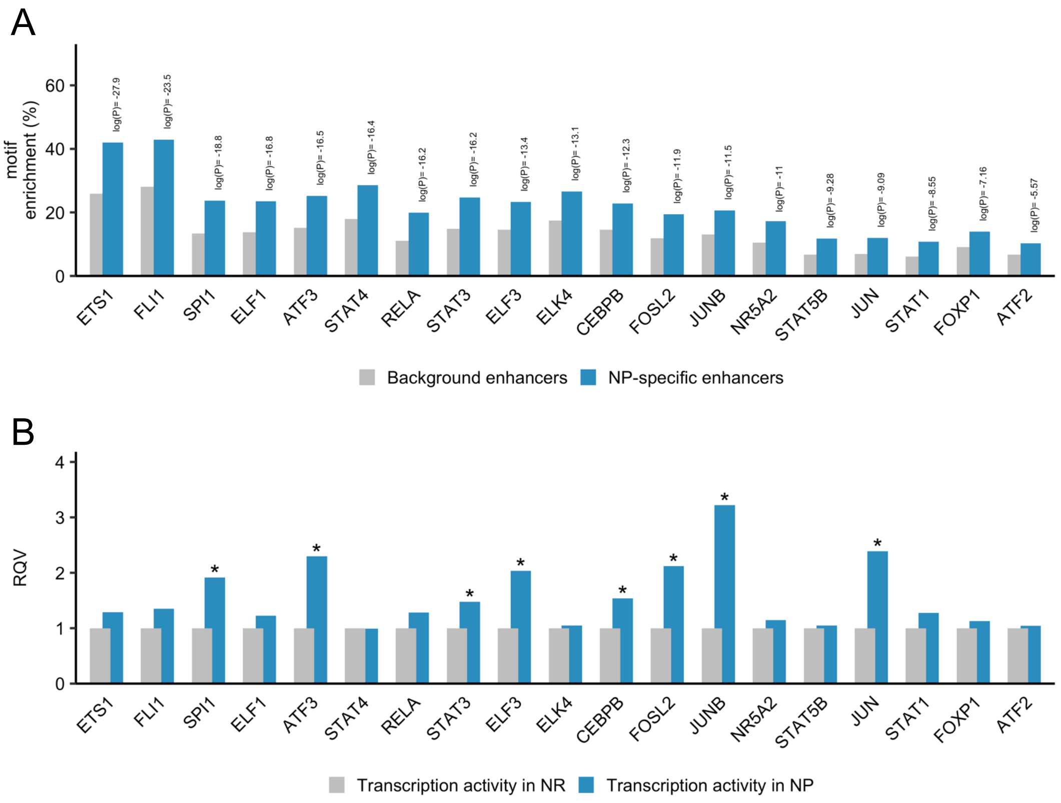


**Supplementary Figure 2.** (A) Complete list of TFs with motifs significantly enriched in NP-specific enhancers ($p < 0.05$, $q < 0.05$, enrichment foldchange > 1.5, target sequences with motif > 10% by HOMER). Only TFs with high transcription levels in NP livers (normalized counts > 500 in NP livers) are shown. (B) Transcription fold-change of TFs highlighted in (A). * = $P < 0.05$, adjusted $P < 0.2$, log2 foldchange < 0 in NR compared to NP (Wald test; DESeq2).

| | NR | NP | P-value |
|---|---|---|---|
| Glycemic status | 1 NG / 3 IGT / 2 T2D | 1 NG/ 4 IGT / 3 T2D | NA |
| rs738409 genotype | 2 CC / 4 CG | 3 CC / 5 CG | NA |
| Gender (Female; Male) | 6 F / 0 M | 8 F / 0 M | NA |
| Age | 48.83 ± 4.54 | 39 ± 9.49 | 0.027 |
| Height (cm) | 156.83 ± 8.70 | 159.44 ± 3.98 | 0.519 |
| Weight (kg) | 125.95 ± 17.56 | 129.66 ± 11.65 | 0.665 |
| BMI | 51.13 ± 5.22 | 50.95 ± 3.54 | 0.943 |
| Waist Cir. (cm) | 138.4 ± 15.34 | 137.75 ± 14.46 | 0.941 |
| Hip Cir. (cm) | 149.4 ± 13.99 | 148.13 ± 8.46 | 0.860 |
| Waist-to-hip ratio | 0.93 ± 0.08 | 0.93 ± 0.09 | 0.980 |
| Hemoglobin A1C | 0.069 ± 0.02 | 0.07 ± 0.01 | 0.812 |
| Fasting plasma glucose (FPG; mM) | 8.75 ± 4.99 | 7.21 ± 1.85 | 0.499 |
| Total Cholesterol (mM) | 4.04 ± 0.84 | 5.06 ± 0.74 | 0.040 |
| HDL Cholesterol (mM) | 1.27 ± 0.15 | 1.27 ± 0.21 | 0.956 |
| LDL Cholesterol (mM) | 2.15 ± 0.76 | 3.09 ± 0.81 | 0.047 |
| Triglyceride (mM) | 1.38 ± 0.57 | 1.53 ± 0.40 | 0.596 |
| Total : HDL cholesterol | 3.24 ± 0.87 | 4.08 ± 0.97 | 0.114 |
| Smoking status<br>(1 = smoker / 2 = non-smoker / 3 = NA) | 0/5/1 | 0/8/0 | 0.363 |
| % steatosis | 13.5 ± 14.54 | 63.13 ± 16.46 | 0.0001 |
| Steatosis grade (0/1/2/3) | 2/3/1/0 | 0/0/4/4 | 0.0014 |
| Ballooning (0/1/2/3) | 6/0/0/0 | 0/6/2/0 | 0.0001 |
| Lobular inflammation (0/1/2/3/4) | 3/3/0/0/0 | 0/3/4/1/0 | 0.0029 |
| Portal inflammation (0/1/2/3/4) | 4/2/0/0/0 | 0/7/1/0/0 | 0.0113 |
| Hepatic fibrosis (0/1/2/3/4) | 5/1/0/0/0 | 0/3/5/0/0 | 0.0001 |
| NAFLD activity score (/11) | 1.5 ± 1.38 | 7 ± 1.51 | 0.000016 |
| Alanine aminotransferase (ALT; U/L) | 24.33 ± 8.29 | 44.88 ± 20.77 | 0.0300 |
| Aspartate aminotransferase (AST; U/L) | 19.67 ± 2.42 | 30.75 ± 7.27 | 0.0030 |
| AST/ALT | 0.86 ± 0.19 | 0.78 ± 0.28 | 0.5148 |
| Gamma-glutamyl transferase<br>(GGT; U/L) | 26 ± 7.07 | 43.13 ± 22.77 | 0.0773 |
| Bilirubin total (umol/L) | 6 ± 1.79 | 9.13 ± 3.44 | 0.0500 |
| Bilirubin direct (umol/L) | 1.83 ± 0.41 | 2.13 ± 0.99 | 0.4699 |
| Alkaline phosphatase (ALP; U/L) | 100 ± 20.81 | 74.38 ± 12.26 | 0.0290 |

**Supplementary Table 1.** Clinical characterization of patients with obesity, stratified into either NP or NR phenotype.

**Supplementary Table 2.** Summary of ChRO-seq mapping statistics and identification of active transcriptional regulatory elements (TREs).

(large data set)

**Supplementary Table 3**. Differential analysis of transcription (ChRO-seq signal) at gene bodies and transcriptional regulatory elements (TREs) in NR compared to NP livers. Genes with $P < 0.05$, adjusted $P < 0.2$, log2 foldchange > 0 (or < 0), average normalized counts > 100 were defined as differentially transcribed genes (Wald test; DESeq2). TREs with $P < 0.05$, adjusted $P < 0.2$, log2 foldchange > 1 (or < -1) were defined as differentially active TREs (Wald test; DESeq2). NASH-resistant (NR), n = 6; NASH-prone (NP), n = 8.

(large data set)

**Supplementary Table 4.** Enhancer density analysis. In this analysis, the number of NP- or NR-specific enhancers within a window of +/-100 kb around the annotated TSS was counted for each transcribed gene (average normalized counts > 100 across all the samples). NP-activated genes associated with NP-specific enhancers and NR-activated genes associated with NR-specific enhancers are shown.

(large data set)

**Supplementary Table 5.** Analysis defining NP and NR-specific enhancer hotspots. Parent: the coordinates of a given stitched enhancer; Rank: the ranking based on the total activity (ChRO-seq signal); Total Signal: the sum of the ChRO-seq signal from each of the individual enhancers in a stitched enhancer; NumTRE: the number of individual enhancers within a stitched enhancer; Length: the length of a stitched enhancer; Super: denotes whether the total signal of a stitched enhancer is strong enough to be defined as an enhancer hotspot; Children: the coordinates of each individual enhancer within a stitched enhancer.

(large data set)

**Supplementary Table 6.** TF cistrome analysis. NP-activated genes associated with NP-specific enhancers that harbor one or more NP-associated TFs (PU.1, ATF3, STAT3, ELF3, AP-1, and CEBPB) are shown. NR-activated genes associated with NR-specific enhancers that harbor one or more NR-associated TFs (HNF4A, HNF1B, PPARA, and FOXA2) are shown.

(large data set)

| Intersection analysis | | | |
|---|---|---|---|
| Trait | Number of SNPs and associated LD buddies | Intersection analysis | |
| | | NP-specific enhancers | NR-specific enhancers |
| NAFLD | 17 | 0 | 0 |
| Liver injury | 102 | 0 | 0 |
| Liver enzymes | 1218 | 0 | 0 |
| Bilirubin | 285 | 0 | 0 |
| BMI | 473 | 0 | 0 |
| CRP | 1033 | 3 | 0 |
| HbA1c | 310 | 0 | 0 |
| Serum albumin | 43 | 0 | 0 |
| Glucose | 548 | 0 | 0 |
| Insulin | 316 | 0 | 0 |
| T2D | 1555 | 0 | 0 |
| Hypertension | 929 | 0 | 0 |
| Cardiovascular disease | 109 | 0 | 0 |
| Triglycerides | 2153 | 0 | 0 |
| HDL cholesterol | 2609 | 0 | 1 |
| LDL cholesterol | 1601 | 0 | 0 |

| Hits of SNP-containing NP-specific enhancers | | | |
|---|---|---|---|
| Tag SNP | Query SNP | R2 | p-value |
| rs1205 | rs7553007 | 0.9485 | 2E-16 |
| rs1205 | rs876537 | 0.8914 | 0.000000001 |
| rs1205 | rs2794520 | 0.9744 | 2E-186 |

| Hit of SNP-containing NR-specific enhancers | | | |
|---|---|---|---|
| Tag SNP | Query SNP | R2 | p-value |
| rs3847302 | rs3890182 | 0.9978 | 3E-10 |

**Supplementary Table 7**. Summary of intersection analysis with disease/trait-associated SNPs and enhancers uniquely activated in human NP or NR livers.

| Motif scan of NP-specific enhancer (chr1:159711980-159712490; hg38) containing rs1205 | | | | | | | |
|---|---|---|---|---|---|---|---|
| motif_id | start | stop | strand | score | p-value | q-value | matched_sequence |
| ZFP82_HUMAN.H11MO.0.C | 451 | 474 | + | 13.5379 | 7.82E-06 | 0.00734 | CTTCTGTCCTCACAGTCTCTCT**C**C |
| MAZ_HUMAN.H11MO.0.A | 454 | 475 | - | 10.4085 | 5.42E-05 | 0.0254 | TGGAGAGAGACTGTGAGGA**C**AG |
| SMAD4_HUMAN.H11MO.0.B | 463 | 475 | + | 11.6837 | 4.51E-05 | 0.0441 | CAGTCTCTCT**C**CA |

| Motif scan of NR-specific enhancer (chr9:104886231-104886650; hg38) containing rs3847302 | | | | | | | |
|---|---|---|---|---|---|---|---|
| motif_id | start | stop | strand | score | p-value | q-value | matched_sequence |
| NANOG_HUMAN.H11MO.0.A | 78 | 94 | - | 13.589 | 8.79E-06 | 0.00335 | TATATTGAAA**T**GCAAAG |
| PO5F1_HUMAN.H11MO.0.A | 78 | 93 | - | 13.4091 | 0.0000113 | 0.00437 | ATATTGAAA**T**GCAAAG |
| HNF4G_HUMAN.H11MO.0.B | 74 | 87 | - | 12.6061 | 0.0000222 | 0.016 | AAA**T**GCAAAGTGCA |
| HNF4A_HUMAN.H11MO.0.A | 74 | 87 | - | 12.0606 | 0.0000268 | 0.0107 | AAA**T**GCAAAGTGCA |

**Supplementary Table 8**. Top: Motif analysis of NP-specific enhancer (chr1:159711980-159712490; hg38) containing rs1205. Bottom: Motif analysis of NR-specific enhancer (chr9:104886231-104886650; hg38) containing rs3847302. Motif scan analyses were performed with MEME suite. Only motifs that overlap with SNPs (bold letters) and have q-value < 0.05 are reported.
